# *SpliceImpactR* maps alternative RNA processing events driving protein functional diversity

**DOI:** 10.1101/2025.06.20.660706

**Authors:** Zachary Peters Wakefield, Ana Fiszbein

## Abstract

Alternative RNA processing is a key regulator of gene expression, driving transcript and proteomic diversity essential for cell function. However, the precise impacts of different alternative RNA processing events on protein function remain poorly understood. Here, we introduce SpliceImpactR, an open-source tool that systematically identifies RNA isoform switches—including alternative first and last exons, exon skipping, intron retention, hybrid exons, and splice site variation—across the human transcriptome, and predicts their impact on encoded proteins. We find that intron retention and hybrid exons frequently alter the coding potential of transcripts. Strikingly, when both isoforms remain protein-coding, 87% of alternative RNA processing events result in substantial changes to the protein sequence. Among these, alternative last exons drive the largest changes, frequently disrupting protein-protein interactions. Across human tissues, alternative first exons drive the largest relative changes in proteins, often resulting in tissue-specific protein domains. Notably, frameshifts introduced by alternative splicing are often rescued by co-regulated alternative downstream exons, suggesting a buffering mechanism in isoform regulation. Moreover, we show that alternative splicing exhibits gradual, tissue-specific variation, rather than binary on/off behavior, enabling conserved regulation of protein domains across tissues. Together, our results provide a proteome-wide view of splicing regulation, uncovering widespread, context-dependent impacts of alternative splicing on the human proteome.

## INTRODUCTION

Early transcriptomic studies showed that most human genes generate multiple mRNA isoforms through alternative splicing of internal exons. More recently, however, alternative first and last exons have been identified as even greater contributors to transcript diversity^1^. Collectively, alternative RNA processing events impact nearly every aspect of gene function, from transcript subcellular localization and translational efficiency to protein domain architecture and post-translational modifications^2–5^. These changes can reshape protein structures, modulate post-translational modification sites, and rewire protein-protein interaction (PPI) networks^5,6^.

There are several major classes of alternative RNA processing events. Skipped exons (SE) are internal exons that are included in some transcripts and excluded in others. Retained introns (RI) are introns that remain in the mature transcript rather than being spliced out. Alternative 3′ and 5′ splice sites (A3SS and A5SS) cause variation at exon boundaries by shifting the precise splice junctions. Mutually exclusive exons (MXE) involve pairs of exons where only one is included in the mature mRNA. Alternative first exons (AFE) are regulated by different promoters, while alternative last exons (ALE) are associated with the use of alternative polyadenylation sites. Within first exons, alternative transcription start sites (TSS) differ in the first nucleotide transcribed. Within last exons, alternative transcription end sites (TES) refer to the use of different polyadenylation sites, leading to alternative 3′ untranslated regions (UTRs). Together, these events dramatically increase isoform complexity and are tightly regulated.

The functional impact of alternative RNA processing in protein-coding genes is particularly pronounced in tissue-specific contexts. Distinct splicing patterns drive the production of specialized protein isoforms, enabling precise regulation of cellular processes^7–10^. For example, neurons exhibit extended 3′ untranslated regions and uniquely spliced exons critical for synaptic function, while immune cells rely on alternative splicing to diversify signaling pathways and adapt rapidly to pathogens^11,12^. Dysregulation of these processes is increasingly linked to disease: in cancer, distinct isoforms can promote tumor progression, metastasis, and immune evasion^13–19^. However, the functional consequences of these changes remain poorly understood, particularly in complex physiological and disease contexts.

In recent years, numerous computational tools have been developed to predict the downstream functional consequences of alternative RNA processing. These tools enable the analysis of alternative splicing changes across various conditions^20^, the prediction of protein domains that are altered^21–24^, and facilitate domain-integrated protein-protein interaction studies^9,24,25^. Additionally, they support the assessment of evolutionary conservation of alternative RNA processing events^26^, and long-read sequencing approaches for comparing protein isoforms^27^, and visualization of how isoform changes may impact transcript and protein sequence^28^. Still, the specific implications of each alternative RNA processing event on the functional proteome and the global consequences in protein function are not resolved.

Alternative RNA processing is not a series of isolated events but a highly coordinated process, where decisions at one stage can significantly influence downstream choices. For example, alternative first and alternative last exon usage are interlinked within a positional initiation-termination axis, where decisions at the 5’ end directly affect those at the 3’ end^29,30^. Coupling also occurs between alternative splicing and 3’ end processing^31^ as well as between alternative first exons and alternatively spliced internal exons^32–34^. In fact, we recently discovered more than 100,000 human *hybrid exons* that can both serve as terminal or internal exons. Hybrid first exons (HFE) can act as first exons or internal exons, while hybrid last exons (HLE) act as internal or last exons in different transcripts^35^. Despite these well-documented interactions among RNA processing events, there are currently no computational pipelines capable of integrating all these aspects to reliably predict the functional impact of isoform-specific alterations at the gene level.

To address this systematically, here we developed SpliceImpactR, an integrative, open-source R package designed to unite exon-level information from RNA-seq data to reveal how differentially used alternative RNA processing events coordinately shape protein function. Running SpliceImpactR across all annotated human isoforms and 17,350 samples from 54 human tissues, we found that retained introns and hybrid exons drive the most significant shift toward non-protein-coding mRNAs, while alternative last exons lead to the largest changes in protein primary sequences and domains, and alternative 5’ and 3’ splice sites introduce the most substantial frameshifts. Interestingly, we found that, across human tissues, alternative first exons drive the largest relative changes in proteins, specifically resulting in tissue-specific protein domains.

## RESULTS

### Alternative RNA processing has widespread impact on protein primary sequences

SpliceImpactR begins by importing all alternative RNA processing events that differ across conditions (Step 1). It leverages rMATS^36^ and the HIT index^35^ to detect changes in alternative RNA processing events, including alternative first exons (AFE), hybrid first exons (HFE), retained introns (RI), skipped exons (SE), alternative 5’ and 3’ splice sites (A5SS, A3SS), mutually exclusive exons (MXE), hybrid last exons (HLE), and alternative last exons (ALE). It then performs differential inclusion analysis to identify condition-specific pairs of transcripts that differ in at least one alternative RNA processing event (Step 2). These differentially included events are mapped to transcript and protein annotations to evaluate their effect on coding potential, sequence similarity, and frameshifts (Step 3). SpliceImpactR then uses InterPro^37^ to annotate domain-level changes (Step 4), assess domain usage shifts across conditions (Step 5), and integrate domain– domain interaction data from 3did^38^ to predict impacts on protein-protein interactions (Step 6). Finally, it conducts a global analysis to detect co-regulated splicing events and quantify the relative contribution of each alternative RNA processing type (Step 7; see Figure 1A).

**Figure 1.**
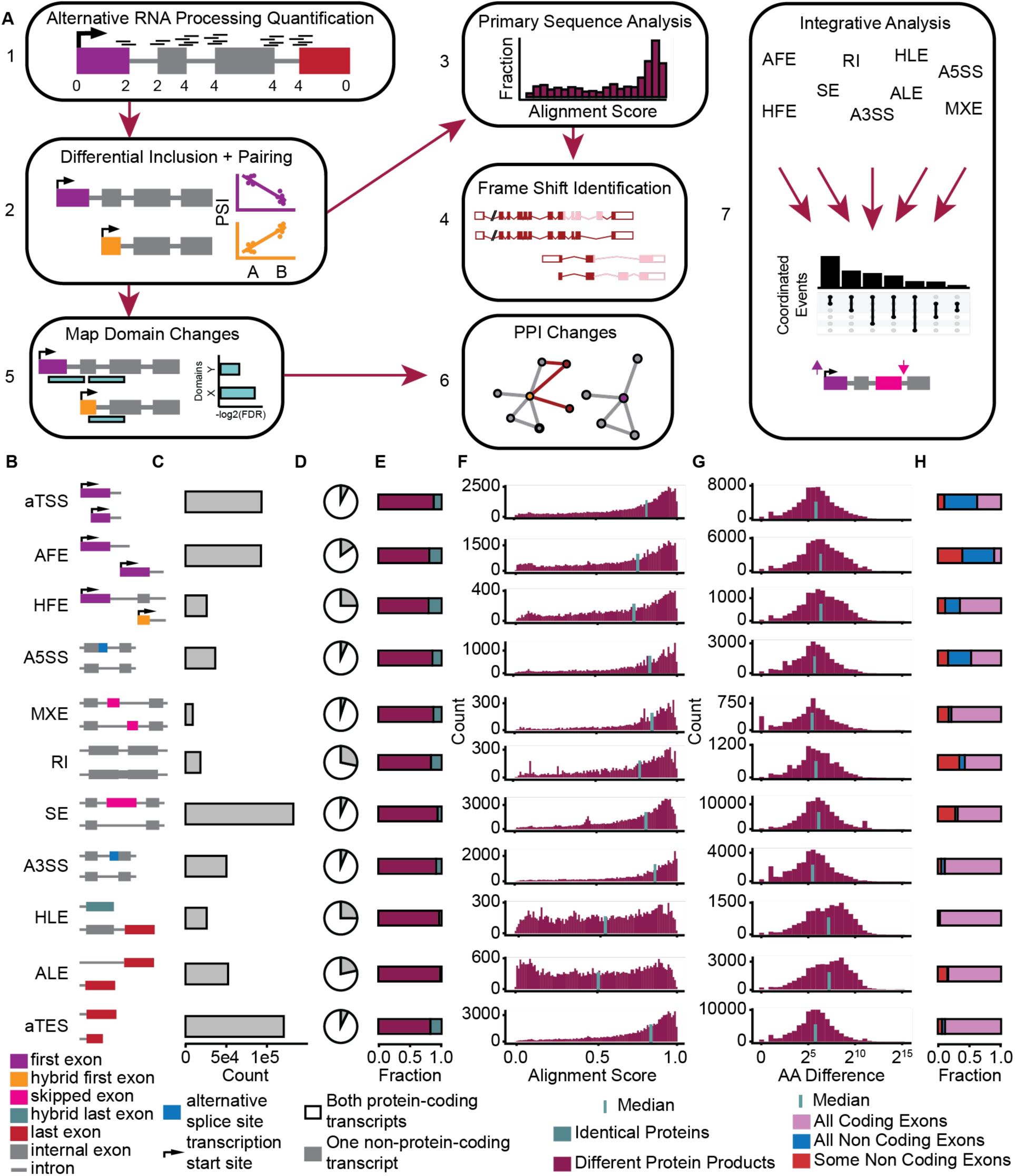
SpliceImpactR and the widespread impact of alternative RNA processing on protein primary sequences. (A) (1) *SpliceImpactR* takes as input processed RNA-seq data derived from the HIT Index and rMATS pipelines. (2) Significant differential RNA processing events are identified based on statistical testing and delta percent-spliced-in (ΔPSI) thresholds. Transcripts differing by at least one such event are grouped into matched pairs. (3) Primary amino acid sequence comparisons are performed to assess sequence similarity and classify transcript-level differences. (4) InterPro domain annotations are mapped to the protein products of these transcript pairs to detect localized domain changes. (5) Identified domain changes are aggregated to reveal global patterns and trends in domain architecture alterations. (6) Known domain-domain interactions from 3did and PPI-DomainMiner databases are used to determine whether alternative RNA processing alters protein–protein interaction interfaces. (7) A comprehensive integrative analysis is performed across all differential events to identify co-regulated splicing patterns and predict their aggregated effects on proteins. (B) Diagrams of the 11 forms of alternative RNA processing considered: aTSS (alternative transcription start sites), AFE (alternative first exons), HFE (hybrid first exons), A5SS (alternative 5’ splice sites), MXE (mutually exclusive exons), RI (retained introns), SE (skipped exons), A3SS (alternative 3’ splice sites), HLE (hybrid last exons), ALE (alternative last exons), and aTES (alternative transcription end sites). (C) Count of transcript pairs that differ in a given event among 171,397 annotated isoforms. (D) Fraction of transcript pairs differing in a given event that change protein-coding status. (E) Fraction of paired transcripts differing in a given event that result in either identical proteins or different proteins, when both transcripts are protein-coding. (F) Distribution of primary sequence similarity between protein isoforms encoded by each transcript differing in a given alternative RNA processing event. Alignment score is calculated through sequence alignment using BLOSUM62 (# matched amino acids / # unmatched amino acids). Vertical line shows the median alignment score. (G) Distribution of changes in protein length (log₂ scale) associated with each alternative RNA processing event type. The vertical line indicates the median length change for each event. (H) Proportion of alternative RNA processing events based on the genomic location of involved exons — classified as entirely within coding regions (“all coding exons”), entirely outside coding regions (“all non-coding exons”), or a mix of coding and non-coding regions (“some non-coding exons”).

We first use SpliceImpactR to explore the full spectrum of protein diversity generated by all forms of alternative RNA processing annotated in the human transcriptome. Using the full set of 171,397 isoforms annotated in the last human genome of reference, we identified all transcript pairs with at least one difference in any of the 11 alternative RNA processing events considered (Figure 1B), resulting in 654,444 paired transcripts. Notably, SEs were the most prevalent alternative processing event, differentially included in 132,100 transcript pairs, followed by aTES, aTSS, and AFEs. In contrast, MXE and RI were the least common, occurring in 8,617 to 18,044 transcript pairs (Figure 1C). We then analyzed which of these alternative processing events completely disrupt the coding capacity of their transcripts. We found that nearly 30% of RIs cause its transcripts to become non-protein coding when retained. Similarly, hybrid exons—whether used as terminal or internal exons—are strongly associated with coding status changes, with 25% of cases shifting from a protein-coding to a non-coding transcript (Figure 1D). Alternative first and last exon usage can also often lead to the loss of protein-coding transcripts. However, SEs, aTSSs, and aTESs very rarely lead to changes in protein-coding potential and both transcript isoforms remain protein-coding in most cases. Importantly, when alternative RNA processing events do not alter the protein-coding status of transcripts, 87% still lead to significant changes in the resulting protein sequences (Figure 1E). Despite not being the most frequent alternative RNA processing events, ALEs and HLE appear to drive the most significant changes in protein sequences with only ∼3% maintaining the identical protein sequence (Figure 1E).

Paired proteins that only differ in the usage of an ALE have the lowest median alignment score (0.51, Figure 1F) and the largest change in protein length, with a median change of 145 amino acids (Figure 1G). One of the most striking examples is the TTN gene, where the use of an upstream ALE shortens the transcript by more than 98 kilobases (Figure S1A). Interestingly, AFEs, and HFEs rank next in driving substantial changes in protein sequences, with median alignment scores of 0.76 and 0.73 and a median change of 78 and 80 amino acids respectively (Figure 1F, G). This indicates that alternative changes at the 3’ end of transcripts have the greatest impact on protein sequence, followed by changes at the 5’ end, while alternative RNA processing events occurring within gene bodies result in smaller changes. Interestingly, all RNA processing events include cases where the alternative event does not alter the final protein product. This is particularly common in AFEs and aTES as more than 50% of both of their variants are located in non-coding regions (Figure 1H). When events occur within coding regions, we primarily observe intuitive cases where changes affect the resulting proteins, such as an AFE in *CHRAC1*, an MXE in *UBE2C*, and an A3SS in *C16orf92* where each form has distinct coding nucleotides (Figure S1B). However, we identified ∼3000 unexpected cases where changes within coding regions do not alter the resulting protein. These are often represented by short identical coding sequences, such as in the AFE of *MRM2* and the RI of *SPTLC1*. In *DBMT1, a* complex genomic rearrangement involving repeating sequences results in MXEs that do not alter the protein sequence (Figure S1C). In these examples, we see an increase in the mRNA diversity, however no increase in proteomic diversity – the redundant isoforms can undergo alternative splicing without altering the final protein.

### Co-regulation of alternative RNA processing rescues coding frame shifts

Alternative RNA processing often leads to significant protein sequence changes by introducing frameshifts within coding regions. This commonly occurs when SE or alternative splice sites include or exclude nucleotides not a multiple of three. For instance, in *CCN5*, an alternative 5’ splice site extends exon 2 by eight nucleotides, shifting the reading frame in exon 4 and producing a completely different protein sequence (Figure 2A). Such frameshifts alter how nucleotides are read, generating different codons, different amino acids, and ultimately completely distinct protein products.

**Figure 2.**
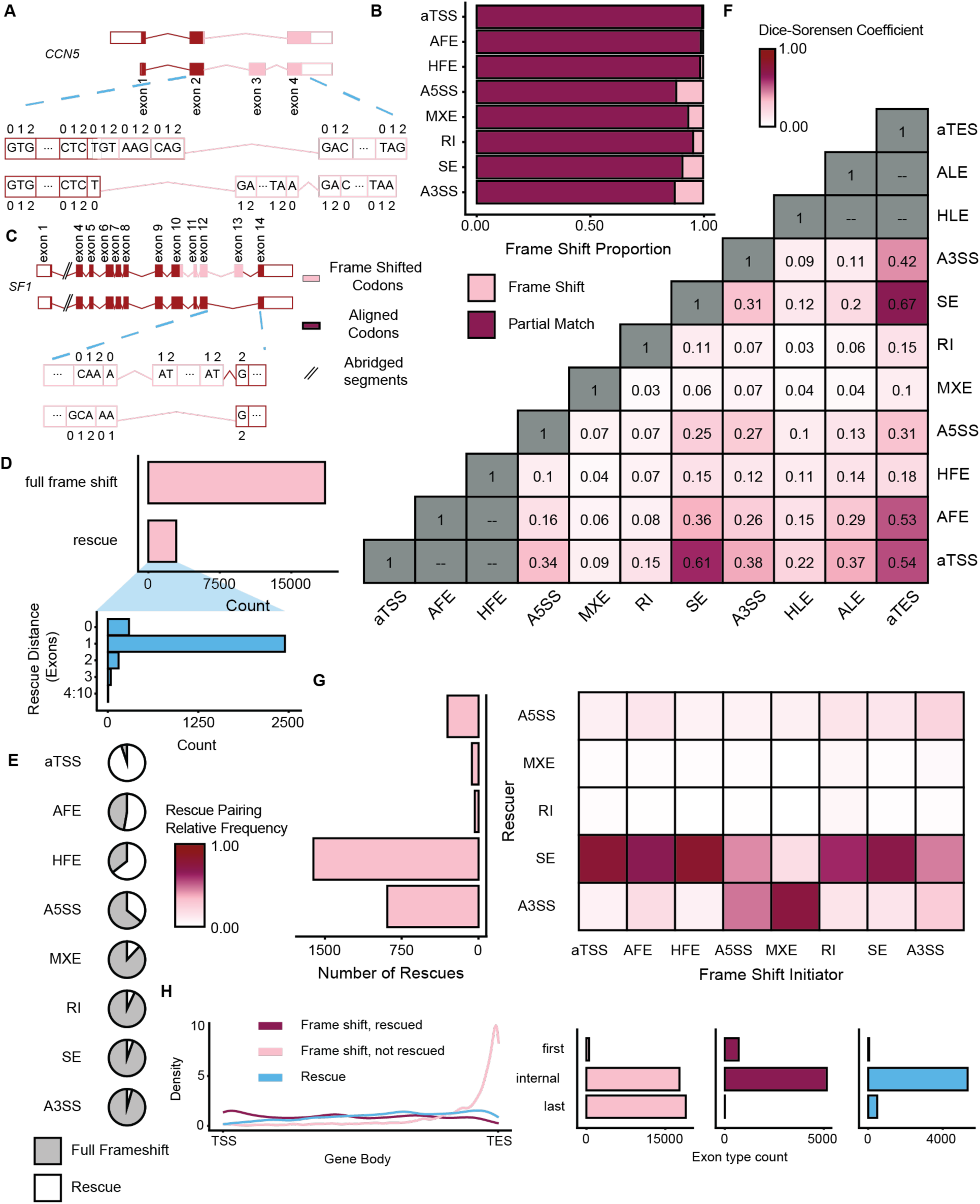
Analysis of frame shift impacts across annotated human protein-coding transcripts. (A) Example from the *CCN5* gene illustrating a frame shift initiated by an alternative 5’ splice site (A5SS) in exon 2, which introduces an 8- base extension at the downstream end of exon 1. This shift propagates through exon 4, altering the reading frame of all downstream overlapping exons. (B) Proportion of protein-coding transcript pairs affected by frame shifts. (C) Example of a rescued frame shift in *SF1*. An A5SS in exon 10 causes a frame shift, which is then restored by the skipping of exon 13 (SE), realigning the downstream reading frame. (D) Total number of observed full frame shifts and frame shift rescues. Inset shows the distribution of rescue distances (in number of exons) between the initiating and rescuing events. (E) Proportion of frame shift–initiating events that are rescued, grouped by event type. (F) Dice-Sørensen coefficient measuring similarity among transcript sets for each alternative RNA processing event. Cases where one event is a subset of another (e.g., HFE within AFE) or perfect overlap are greyed out. (G) (Left) Count of rescue events grouped by alternative RNA processing type. (Right) Heatmap showing the proportion of rescuing events corresponding to each type of initiating frame shift event (normalized by column). (H) (Left) Density plots showing the distribution along the gene body of initiating frame shift events that are rescued, rescuing events, and unrescued frame shifts. (Right) Categorization of event-associated exons as first, internal, or last exons. This categorization refers to the exon type where the first frame shifted bases are seen in the transcript pairs.

We found these shifts mainly stem from internal splicing events, with alternative 3’ and 5’ splice sites accounting for frameshifts in roughly 14% of transcript pairs (Figure 2B). Interestingly, we observed thousands of cases where an alternative processing event induces a frameshift that is corrected by a downstream event. For instance, in SF1, exon 10’s alternative 5’ splice site initiates a frameshift that continues across three exons until an SE event at exon 13 realigns the reading frame for the remaining exons (Figure 2C). This type of event effectively replaces a whole section of the protein with a different amino acid sequence while preserving the terminal ends of the protein. Notably, we observed frame rescues in approximately 13.6% of all frameshift cases with almost 10% of them rescued within the same exon (Figure 2D). An example is TRAPPC2L, where an aTSS induces a frameshift, and an A5SS in the same exon restores the reading frame (Figure S2A). Beyond same-exon rescues, most occur in the following exon, but some corrections are observed up to 10 exons downstream (Figure 2D). Interestingly, frameshifts arising from alternative initiation events, such as aTSS, AFE, and HFE, are largely corrected, whereas those resulting from internal splicing events often remain unrescued (Figure 2E).

To investigate which alternative RNA processing events commonly rescue others, we first analyzed patterns of co-occurrence across transcript pairs. We found that transcripts differing by at least one RNA processing event typically differ by three, indicating frequent combinatorial regulation (Figure S2B). Strong associations emerged between internal splicing events and alternative transcript starts and ends (Figure 2F). This is consistent with previous work showing functional coupling between alternative first and internal or last exons^29,30,33^. In fact, the most common combinations involved sTSS, SEs, and aTES (Figure S2C). Given these patterns, it is not surprising that SEs are the most frequent rescuers of frameshifts, often compensating for disruptions introduced by aTSS, AFEs, or HFE (Figure 2G). Mapping frameshifts and rescues along the gene body show that internal exon frameshifts decline towards the 3’ end of genes, while rescue events increase (Figure 2H). However, a significant proportion of unrescued frameshifts are found toward the end of the gene body, with approximately half occurring in the last exon of a transcript (Figure 2H). Many of these cases involve alternative processing events introducing a frameshift only in the final exon— such as in *PIGV*, where an A3SS extends the final exon by 11 nucleotides (Figure S2D).

### Functional domain changes rewire protein-protein interactions

To determine which alternative RNA processing events directly impact protein function, we focused on events where both transcript variants remain protein-coding. Using the R package biomaRt^39,40^, we identified protein domains for all transcript pairs, mapping InterPro domains to each peptide sequence, and identified cases where an alternative splicing event directly resulted in the addition or removal of a domain. Notably, we found that all types of alternative RNA processing events influence protein domain composition to some extent. ALEs have the largest impact on protein domains, with more than 21% directly resulting in changes to functional protein domain content (Figure 3A). In contrast, aTSS events have the lowest impact, with only ∼1.7% affecting a protein domain (Figure 3A). Generally, when an alternative RNA processing event alters protein domains, it affects only one. However, instances exist across all event types where multiple domains are altered (Figure 3B), with the highest recorded case involving changes in up to seven domains. For example, the use of a distal alternative 5’ splice site in exon 4 of the *MSH6* gene results in the inclusion of three additional protein domains essential for DNA mismatch repair (Figure S3A).

**Figure 3.**
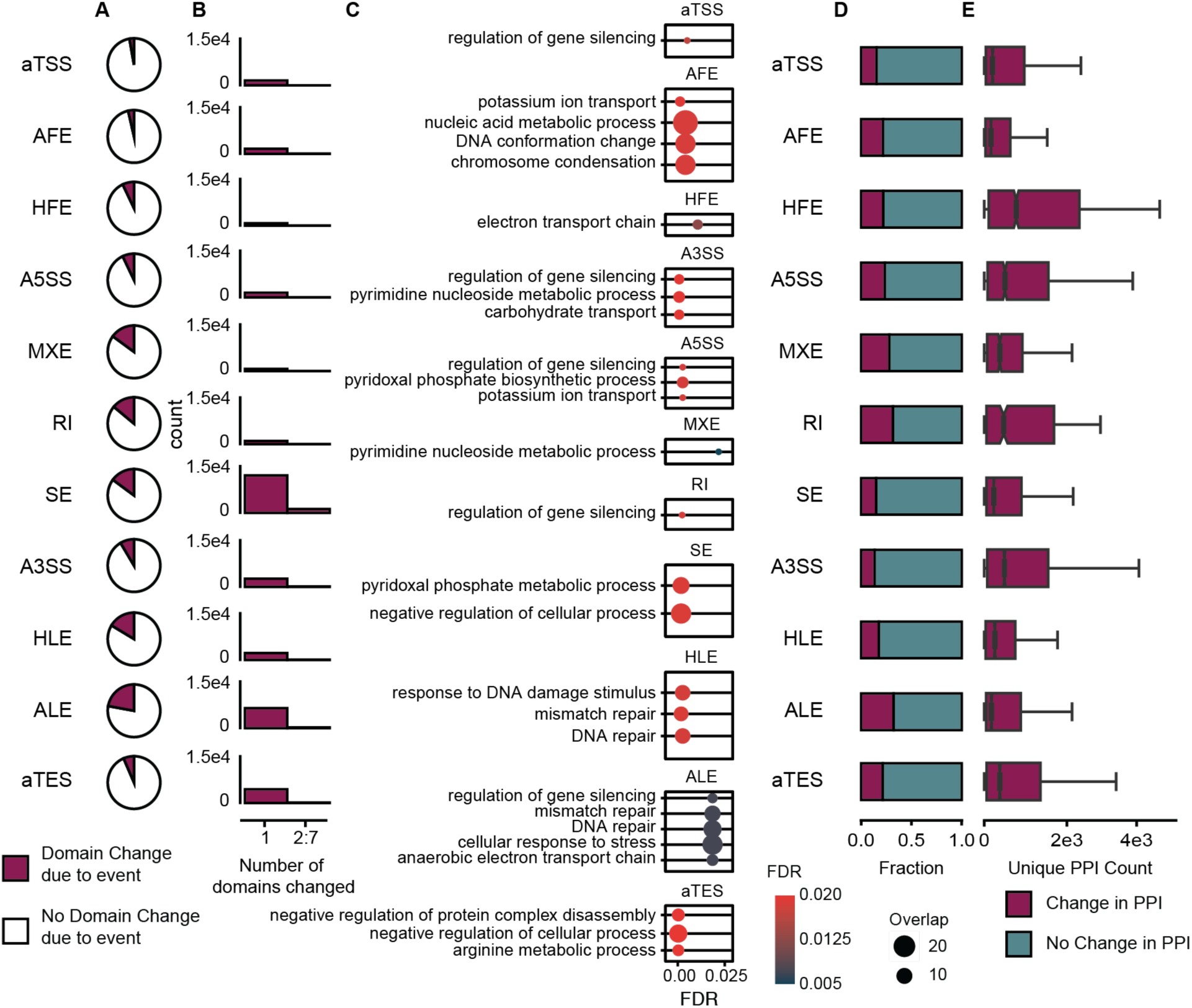
Functional domain changes induced by alternative RNA processing rewire protein–protein interaction networks. **(A)** Proportion of transcript pairs for each alternative RNA processing event type that exhibit changes in InterPro-defined protein domains directly resulting from the alternative event. Domain changes are identified by overlapping the genomic coordinates of alternative events with annotated domain regions. **(B)** Number of domain changes per transcript pair for each event type, categorized as either a single domain change or two or more domain changes. **(C)** Gene Ontology Biological Process enrichment of protein domains altered by each type of alternative RNA processing event, computed using the dcEnrichment() function from the dcGOR R package. Terms with FDR < 0.05 are considered significantly enriched. **(D)** Proportion of transcript pairs with each alternative RNA processing event type that result in rewiring of protein–protein interaction (PPI) networks. Rewiring is defined as the gain or loss of at least one domain–domain interaction, based on the 3did database and domain presence in all human isoforms. **(E)** Distribution of the number of novel interactors gained per isoform within transcript pairs that exhibit PPI rewiring.

Interestingly, protein domains affected by alternative RNA processing are predominantly linked to gene silencing regulation (Figure 3C). AFE-driven domain changes are uniquely enriched for functions related to DNA conformation and chromosome condensation, while those arising from HLE and ALE events are specifically associated with DNA repair (Figure 3C). When analyzing the domain composition across all isoform-altered proteins, we found frequent disruptions in zinc finger domains—particularly C2H2-type and RING-type—which are known to have a wide array of functions including mediation of protein, RNA, and DNA interactions^41^ (Figure S3B). This pattern suggests that alternative RNA processing may preferentially remodel interaction interfaces between proteins. To investigate this, we leveraged 3did’s domain-domain interaction database to predict how such domain alterations could rewire protein-protein interaction networks. Among all event types, ALE and RI change the greatest proportion of protein interaction networks, with 32% of ALE changing at least 1 interactor (Figure 3D). However, protein-protein interaction networks rewired by HFE result in the largest change of novel interactors, with a median addition of 422 interactors per rewired network (Figure 3E). This indicates that alternative RNA processing events significantly alter protein domain composition, and protein-protein interactions with ALE driving the larges changes.

Taken together, our results highlight distinct roles for different alternative RNA processing events in shaping protein diversity. RI are the main contributors to non-coding isoform production, whereas ALE and HLE events drive the most substantial alterations in protein sequences. In contrast, SE, though the most common splicing event, have a relatively minor impact on protein diversity. Notably, ALE events also stand out as major drivers of changes in protein-protein interaction networks.

### Alternative RNA processing has widespread functional consequences across human tissues

To explore how the proteome-level impacts of alternative RNA processing shape tissue identity, we analyzed 17,350 samples from 54 human tissues using data from the Genotype-Tissue Expression (GTEx) project. Using SpliceImpactR in a pairwise fashion, we identified functionally relevant RNA processing changes across tissues. We found that 44% of annotated alternative RNA processing events exhibit significant changes across the human tissues analyzed, indicating that the genome actively leverages a substantial portion of its splicing potential to drive tissue-specific regulation (Figure 4A). Among these tissue-specific events, the median absolute change in percent spliced-in (ΔPSI) was 0.24, with just 1.2% exceeding a ΔPSI of 0.9 across tissues (Figure 4A). These findings indicate that alternative RNA processing is rarely governed by binary on/off switches but instead exhibits a graded, fine-tuned regulation across tissues. Further, we found that 97% of significant changes in AFE and ALE usage involved simple paired switches between two terminal exons, rather than more complex patterns involving three or more exons (Figure 4B). We next clustered human tissues based on the number of significantly differential alternative RNA processing events across nine event types, excluding aTSS and aTES, which are difficult to robustly detect using RNA-seq alone. As anticipated, brain tissues clustered closely and were distinct from other tissue types. Similarly, tissues with related developmental or functional origins—such as blood, cervix, and heart—also grouped together (Figure 4C). These results confirm that tissues with shared characteristics exhibit similar isoform usage patterns and demonstrate that SpliceImpactR effectively captures biologically meaningful alternative RNA processing variation across tissues.

**Figure 4.**
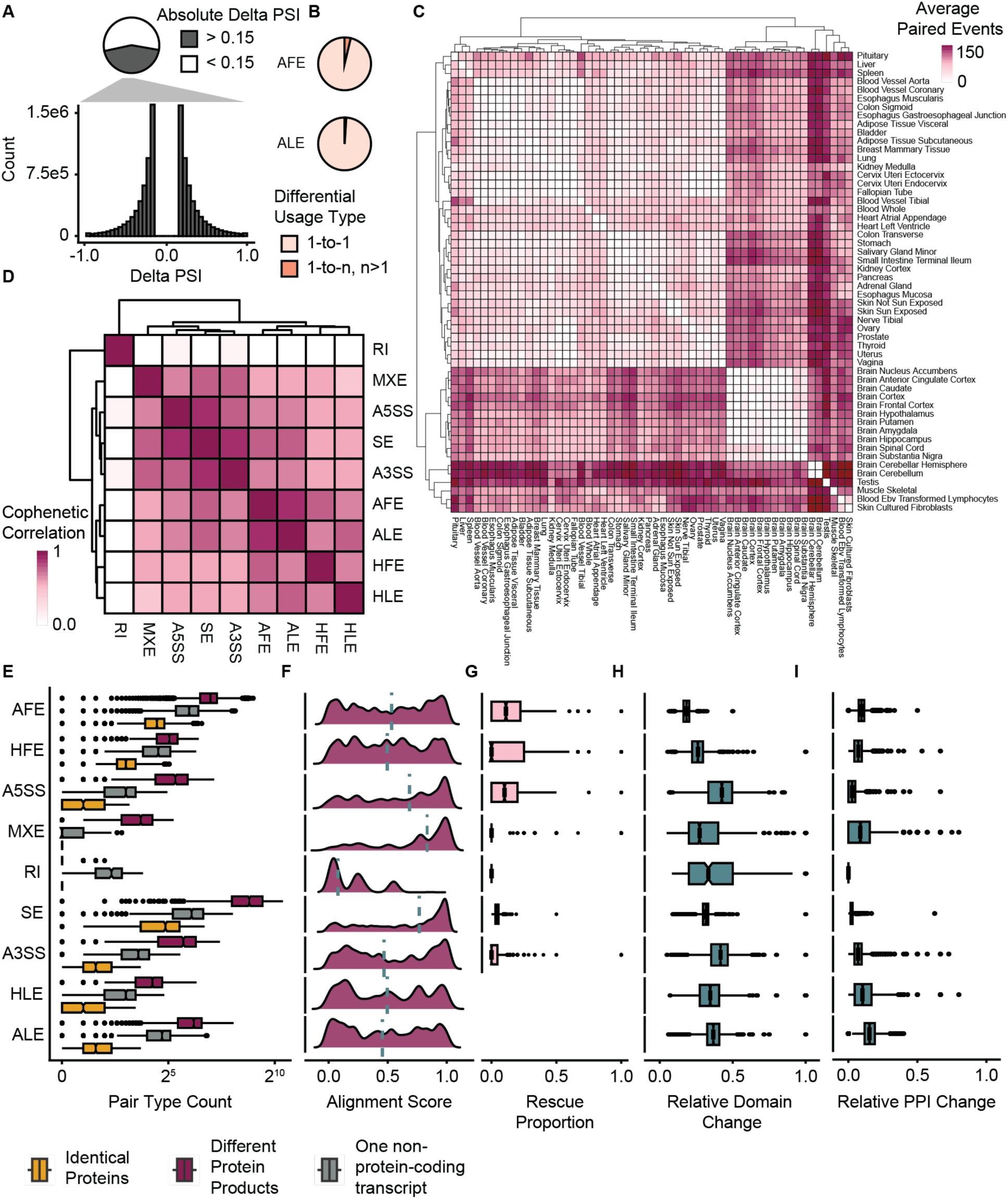
Alternative RNA processing has widespread functional consequences across human tissues. (A) *Top:* Proportion of alternative RNA processing events exceeding the significance threshold (|ΔPSI| > 0.15) across all GTEx tissue pairwise comparisons. *Bottom:* Distribution of ΔPSI values within the significant subset. Values were computed using pairwise runs of SpliceImpactR across GTEx tissues. (B) Proportion of binary 1-to-1 changes (e.g., AFE and ALE) versus more complex patterns (e.g., multiple alternative terminal exons). A 1-to-1 change refers to a binary switch in exon usage across tissues, as opposed to isoform shifts involving three or more variants. (C) Hierarchical clustering (UPGMA) of the mean count of significantly changing events (|ΔPSI| > 0.15, FDR < 0.05) where both isoforms are protein-coding, calculated between each tissue pair using SpliceImpactR. (D) UPGMA clustering based on cophenetic correlation between event-specific clusters. Clusters were derived by computing the Jaccard index for genes undergoing significant, protein-coding isoform switches across tissue pairs for each RNA processing event. (E) Counts of protein-coding status combinations (e.g., coding-to-coding, coding-to-noncoding) across significantly changing events. (F) Distribution of primary protein sequence similarity between isoforms from each event. Alignment scores were calculated using BLOSUM62 as (# matched amino acids / # unmatched amino acids). Dashed line indicates the median score per event type. (G) Proportion of observed frame shifts that are rescued by downstream events. (H) Proportion of events resulting in changes to InterPro domains. A domain is considered “changed” if it overlaps the alternative region and is present in only one isoform. (I) Proportion of events resulting in rewired protein–protein interaction (PPI) networks. Rewiring is defined as any change in predicted interactors based on domain presence/absence, using the 3did domain–domain interaction dataset.

To study how different types of alternative RNA processing are coordinated, we focused on genes with protein-altering events across tissues. We measured tissue similarity using Jaccard indices and hierarchical clustering, which revealed distinct patterns of tissue-specific regulation. Most RNA processing types were co-regulated across tissues, except for intron retention (RI), which behaved independently (Figure 4D). Correlation values among other processing types ranged from 0.55 to 0.87, showing strong coordination. Events affecting transcript ends, including AFE, ALEs and hybrid exons, clustered closely together and separately from internal events such as A5SS, A3SS, and SE. As seen across annotated isoforms, SEs show a much higher correlation of ∼0.76 with AFE and ALE than the median intergroup correlation across tissues (Figure 4D). This indicates that the 3’ and 5’ ends of transcripts are largely co-regulated with SE across human tissues, suggesting that this coordinated regulation plays a key role in shaping tissue-specific transcriptomic identities. Similar to our findings from annotated genome-wide transcript comparisons, RIs primarily influenced protein-coding status, while significant changes in ALEs, A5SS and HLEs usage across tissues rarely resulted in identical proteins (Figure 4E). Among all event types, SE contributed the highest number of significantly changing events that altered amino acid sequences. However, when both transcript variants remained protein-coding, HFEs and RIs led to the most pronounced changes in protein sequence similarity. The average similarity score between protein pairs differing by at least one processing event across tissues was ∼0.50—much lower than the ∼0.75 seen in genome-wide comparisons (Figure 4F). This suggests that although most annotated alternative processing events are not widely used across tissues, the human transcriptome selectively employs those that drive the most substantial protein sequence changes to establish tissue-specific expression.

Interestingly, we found frameshifts and rescues across all significantly changing event types in human tissues, except for RI where no rescues were identified. AFE had the highest proportion of rescued frameshifts, with a median rescue rate of just over 11% (Figure 4G). SE caused the highest absolute number of frameshifts, averaging 115 per tissue, though less than 5% were rescued. Among the genes most changing across all event types, TPM1 showed significant changes in nearly 3000 comparisons (from the total of 12789 comparisons; 1431 pairwise comparisons across 9 alternative RNA processing events), consistent with its known role in tissue-specific function^42,43^. Many other top changing genes also have well-documented roles in tissue-specific isoform expression and alternative splicing (Figure S4A)^44–48^. Genes showing significant tissue-specific changes in alternative RNA processing are enriched in processes involving localization, cell morphogenesis, cytoskeleton organization, and cell growth (Figure S4B). Interestingly, alternative splice site events result in the largest median changes to protein domains (Figure 4H), with RNA recognition motifs among the most frequently affected (Figure S4C). Notably, C2H2-type zinc finger domains were uniquely associated with changes in ALE usage—consistent with our observation that ALEs drive the most substantial rewiring of protein-protein interaction networks across tissues (Figure 4I).

### Functional consequences of alternative splicing in Cerebellum reveals unique tissue-specific behavior

Although SE account for the highest absolute number of protein sequence changes, we found that AFE are proportionally more likely to result in protein-altering outcomes relative to the total number of significantly changing transcript pairs across human tissues (Figure 5A). This suggests that AFE are the most efficiently used mechanism for altering protein sequences across tissues, showing the strongest impact in 47% of all tissue pair comparisons. While all alternative RNA processing types exhibited tissue-specific clustering—particularly between heart, brain, and blood—AFEs displayed the most distinct and robust separation of tissue subtypes (Figure 5B, Figure S5A). Looking at AFE changes in different areas of the human brain, we found a significant shift towards inclusion of proximal AFEs in cerebellum (Figure 5C). We found that 17% of the AFE preferred in cerebellum resulted in a change in inclusion of nuclear localization signal (NLS) motifs – relative to 6% of annotated AFE causing shifts in NLS motifs (Figure 5D). These proximal AFEs are associated with inclusion of specific protein domains, such as cyclin N-terminal and WD40 domains—both of which are well-established regulators of cerebellar differentiation, proliferation, and development (Figure 5E,F)^49–51^. A similar pattern emerged in whole blood, where AFE-driven inclusion of domains known to be critical for blood cell maturation and function was consistently observed across comparisons (Figure S5B, S5C)^52,53^.

**Figure 5.**
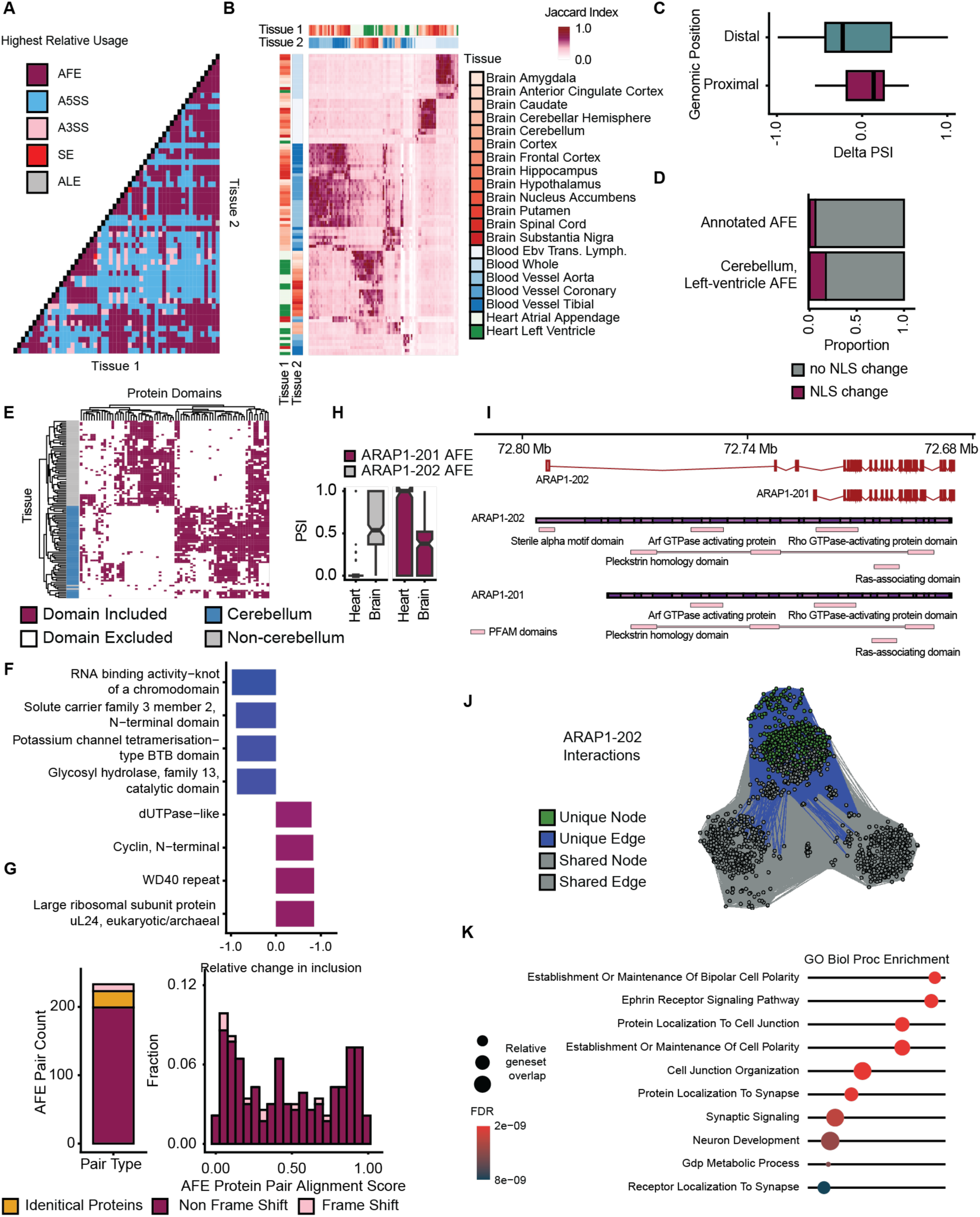
Functional consequences of alternative splicing in the cerebellum reveal distinct tissue-specific signatures. **A.** Heatmap showing the proportion of significantly changing events across tissues for each alternative RNA processing type, calculated as (# significant changes / total observed events). **(B)** UPGMA hierarchical clustering based on Jaccard indices of overlapping genes with significantly changed AFE events (|ΔPSI| > 0.15, FDR < 0.05) across blood, brain, and heart tissues. **(C)** Genomic distribution of cerebellum-enriched AFEs relative to left-ventricle-enriched AFEs. **(D)** Nuclear localization signals (NLS) were identified using 17 consensus motifs from Juane et al. (2021) and scanned using FIMO (MEME-suite). Transcript pairs with significant NLS motif changes (q < 0.05) were marked as showing altered NLS presence. **(E)** Hierarchical clustering (UPGMA) of commonly regulated protein domains associated with cerebellum- preferred versus non-cerebellum-preferred isoforms. Domains shown are those above the median frequency. **(F)** Top included and excluded domains in cerebellum, calculated as (inclusion count / total cerebellum) − (exclusion count / total non-cerebellum). **(G)** (Left) Counts of differentially included AFE events by transcript coding category. (Right) Protein sequence similarity between cerebellum and left-ventricle AFE transcript pairs. **(H)** PSI values of AFEs in the ARAP1 gene across heart and brain tissues. **(I)** (Top) Transcripts associated with each AFE in ARAP1; (Bottom) InterPro domain architecture of proteins encoded by these transcripts. **(J)** Changes in protein–protein interaction networks between ARAP1- 202 and ARAP1-201 driven by domain differences. **(K)** GO Biological Process enrichment of unique protein interactions enabled by the additional domain in ARAP1-202, generated using the hypeR R package.

Many AFEs across blood, brain, and heart shift transcripts between protein-coding and non-coding forms, while most produce distinct protein isoforms (Figure 5G). A striking example is the ARAP1 gene, where a shift between an upstream and a downstream AFE is observed in heart versus brain tissues (Figure 5H). The two resulting transcripts primarily differ in their three most upstream exons, leading to functional differences in the proteins they encode. The cerebellum isoform includes a SAM domain that changes how the protein interacts with others (Figure 5I), and these interactions are enriched in neuron-related processes like brain development and synapse organization (Figure 5J, K), highlighting the role of AFE-driven isoform diversity in neural functions Similar patterns are seen with other processing types—for example, in the *SPEG* gene, which has known heart- and brain-specific isoforms. The left ventricle uses a proximal ALE, while the cerebellum prefers a distal one (Figure S5D). The brain isoform includes a protein kinase domain (Figure S5E), which may enhance its ability to interact with other proteins. These brain- specific interactions are linked to long-term potentiation, a key process in learning and memory (Figure S5F), showing how ALE-driven isoform diversity contributes to neural signaling.

Together, our results suggest that different types of alternative RNA processing events play distinct roles in shaping protein diversity across human tissues. AFE and ALE are not only the strongest drivers of isoform differences but are also associated with distinct SE inclusion patterns across tissues. AFE leads to the largest relative changes in protein sequence and has the highest rate of rescued frameshifts, while ALE drives the most extensive rewiring of protein interaction networks. Notably, specific proximal AFEs appear to play key roles in cerebellar function. These findings highlight how tissue-specific splicing strategically leverages different RNA processing events to remodel the functional proteome.

## DISCUSSION

SpliceImpactR extends beyond traditional analysis methods by integrating holistic, multi-event analyses to uncover interdependencies and co-regulation among alternative RNA processing events. Unlike previously developed tools such as DIGGER, tappAS, and IsoformSwitchAnalyzeR^24–26,28^, SpliceImpactR offers a streamlined, end-to-end framework for downstream analysis following short-read RNA-seq processing with HIT Index and rMATS^35,36^. It also complements other tools—such as the long-read-focused Biosurfer—for comprehensive profiling of isoform diversity^27^ by identifying functional consequences of significantly regulated RNA processing events. Moreover, SpliceImpactR uniquely captures the effects of AFE and ALE – the two forms of alternative RNA processing we found to have the most substantial impact. In contrast to prior studies focused on isolated events^7,9^, SpliceImpactR reveals patterns of co-regulation across event types and provides a global view of how each RNA processing event type contributes to proteomic diversity. Our large-scale tissue-level analysis using GTEx further demonstrates the coordinated regulation of terminal and internal exon events, underscoring its utility in uncovering biologically meaningful splicing patterns. In line with previous transcriptomic studies, we confirm that skipped exons (SE) are the most frequent alternative RNA processing event across human transcripts^7,55,56^. However, despite their abundance, SEs have a comparatively lower impact on protein diversity. Instead, terminal events exert the most substantial influence on protein sequences, supporting earlier findings that terminal splicing events significantly affect proteomic outcomes^5,6,9,10^. We also identify retained introns (RI) as the primary contributors to shifts in protein-coding potential. Beyond individual events, our results highlight extensive co-regulation across splicing types. We observe coordinated usage between internal and terminal events, particularly between SE and both ALE and AFE, echoing previous reports of coupling between internal splicing and alternative promoter usage. Notably, tissue-level comparisons further reveal consistent associations between first and last exon events, reinforcing the spatial coordination of terminal exons^31^.

A key discovery of SpliceImpactR involves patterns of frameshift induction and compensatory rescue, expanding on the previously described “snapback” phenomenon identified by Biosurfer^27^. We show that many frame-disrupting events—especially those at the 5’ end—are subsequently corrected by SEs downstream, preserving protein functionality. This highlights an underappreciated role for SEs in mitigating potentially deleterious consequences of upstream RNA processing variation and supports a model where distinct alternative events cooperate to maintain proteome integrity.

Our analysis shows that more than 44% of annotated alternative RNA processing events are significantly regulated across the 54 human tissues in GTEx. Compared to the full set of annotated events, these tissue-regulated isoforms drive markedly greater changes in protein primary sequence, suggesting a selective regulatory logic that favors isoform usage with the strongest impact on protein structure and function. Among these, AFEs are proportionally more likely to result in protein-altering outcomes relative to the total number of AFE present in human tissues. This suggests that AFEs may function as a key regulatory axis in tissue-specific gene expression. We found changing the 3’ end of a transcript has the greatest possibility to change proteins and rewire interactions in the global set of possibilities in annotations, however in human tissue, we observe that AFE are more often utilized to drive changes.

Domain-level analyses further emphasize the functional consequences of RNA processing, particularly through ALE-mediated changes. Consistent with prior reports on alternative splicing and protein-protein interaction rewiring^4,5,10,24,25^, we observed that ALEs frequently alter zinc finger domains, while tissue-specific contexts show recurrent inclusion of RNA recognition motifs. In specific tissues, we observed a highly conserved number of protein domains included and excluded due to changing alternative RNA processing events. In cerebellum we identified the consistent inclusion of domains such as WD40 repeats and cyclin N-terminal, both of which are strongly associated with neuronal development and cerebellar function^49,50^, reinforcing the idea that terminal exon usage serves a critical regulatory role in shaping cell-type-specific proteomes. Notably, the coordinated use of AFEs and ALEs in genes like ARAP1 and SPEG between cerebellum and heart tissues provides clear examples of how first and last exon choices drive distinct functional outcomes. In these cases, alternative RNA processing choices determine domain presence, protein interaction potential, and tissue-specific roles.

One consideration for SpliceImpactR is its reliance on short-read RNA sequencing data, which may limit the resolution of full-length isoform reconstruction compared to long-read methodologies. However, SpliceImpactR is well-suited for integration with long-read approaches such as Biosurfer, providing a paired long-read and short-read sequencing analysis approach to confirm differences due to alternative RNA processing^27^. Future efforts should focus on experimentally validating predicted changes in protein domains and interaction networks to confirm the functional impact of observed isoform diversity. Additionally, expanding SpliceImpactR to include features like motif enrichment, post-translational modification prediction, and localization signal analysis could further refine the functional interpretation of isoform changes. Applying the framework to clinical datasets may also reveal how alternative RNA processing contributes to disease mechanisms—particularly in cancer and neurological disorders, where splicing dysregulation is a well-established hallmark^13–19^.

## METHODS

### Alternative RNA processing identification and quantification

The hybrid-internal-terminal (HIT) Index^35^ and multivariate analysis of transcript splicing (rMATS)^36^ are applied to short-read RNA-sequencing data to produce relative inclusion values for alternative RNA processing events. The relative inclusion value generated by these tools is a percent-spliced in (PSI) value, calculated through relative junction and exon reads. PSI values are the reads assigned to the specific event (junction and/or exon-covering reads) relative to the other terminal exons or the exclusion form. This metric is calculated differently for rMATS and the HIT

Index respectively, though represents a very similar measure. A generalized form is as such for the PSI of exon *i* in gene *g*:

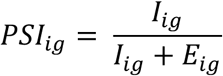

where 𝐼*_ig_* represents the number of reads associated with the inclusion of exon *i* from gene *g*. 𝐸*_ig_* represents the number of reads associated with the exclusion of exon *i* in gene *g*. These tools enabled the identification of alternative events such as AFE, A5SS, MXE, RI, SE, A3SS, and ALE. SpliceImpactR uses the PSI values to generate profiles for samples from each condition. The HIT index also classifies exons -- allowing for the analysis of hybrid first and last exons (HFE, HLE). Events from these tools are imported into R and collapsed to prepare for downstream analysis.

### Condition comparison

Global RNA processing profiles of samples across conditions are compared using event counts and PSI values from HIT Index and rMATS. Event counts are normalized using the mean depth of each event in each sample to control for variations in sequencing and filtered for low expressed events. SpliceImpactR performs comparisons such as event count, distribution of event per gene, and cumulative distribution function of global PSI values are performed to identify shifts in macro splicing patters across conditions. We use a Wilcoxon rank sum test to obtain a p-value for these count changes across condition. We also generate visualizations and output statistics of how HIT Index values assigned to exons are changing – also extracting exons which are significantly changing using the Wilcoxon rank sum test followed by adjusting p-values to FDR.

### Differential analysis and pairing

Differential inclusion of alternative RNA processing is identified through the difference in average PSI value across experimental condition – or a delta PSI score. The pipeline excludes low-coverage events (defaulting to a threshold of 10 inclusion reads for each event) and those with a high number of missing values before applying a statistical model. Next, quality filtering is performed. In the cases of AFE, HFE, HLE, and ALE only exons that are annotated as first or last exons in at least one transcript are included for analysis. PSI values are recalculated using the updated filtered event information. After all filtering is done, the pipeline fills in information for events which are not present in all samples through assigning counts of inclusion of 0 and exclusion of the mean exclusion reads per event per sample. We then remove information relating to genes with solely psi values of 0 on a sample basis. Inclusion reads are extrapolated for AFE, ALE, HFE, HLE from the difference in upstream and downstream junction reads for each exon. Exclusion reads for the same events are further calculated through the sum of the inclusion reads from all exons other than the given exon in each gene.

We implement four statistical models for identifying differential usage of alternative RNA processing events.

1. The first method is exclusively implemented for downstream analysis of the HIT Index. For this, we must first calculate size factors in each sample. This is done using the same method as Anders & Huber (2012)^57^:

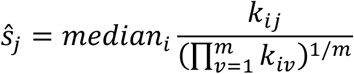

where 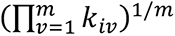 is the geometric mean of reads covering exon *i* in all *m* samples and 𝑘*_ij_* are

the reads covering exon *i* in sample *j*. One difference here is that the reads incorporated into the size factors stem from only the RNA processing event being analyzed. First and last exon processing events use junction-covering reads to calculate size factors instead of exon-covering reads. Similarly, we adapt the method from the updated implementation of Anders et al. (2012) to model negative binomial-distributed reads using a generalized linear model. This model integrates the influence of the sample, exon, and the experimental condition effect on the target exon. To account for different read coverages across samples, we incorporate an offset term based on the previously calculated sample size factors. The negative binomial regression model for the number of reads covering exon *i* in sample *j* (𝑛*_ij_*) is specified as:

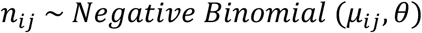

where 𝜇*_ij_* is the expected number of reads covering exon *i* in sample *j*, and 𝜃 is the dispersion parameter. The model used is:

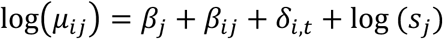

Where 𝛽*_j_* accounts for sample *j*-specific variability. 𝛽*_ij_* captures the effect of exon *i* in sample *j*.

𝛿*_i,t_* represents the interaction between the experimental condition *t* and target exon *i*. The final aspect serves as an offset term, with 𝑠*_j_* being the size factor for sample *j.* The negative binomial regression was implemented using the glm.nb() function from the **MASS** package in R^58^, following the standard workflow to estimate dispersion parameters.

2. We extend this in our second statistical approach to account for the overrepresentation of zeros relative to conventional negative binomial distributions in alternative RNA processing data. We employ a zero-inflated negative binomial (ZINB) model. The excess zeros arise when certain AFE and ALE appear in some samples but not in others, requiring the “missing” exons to be recorded as zero reads and PSI values. Under a standard negative binomial distribution, these additional zeros lead to poor model fit, because of the observation of more zeros than expected. The ZINB model accommodates this by modeling counts as negative binomial and excess zeros as logit:

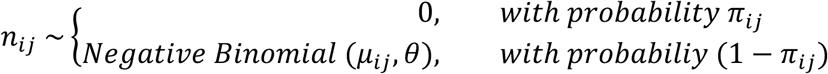

We use the same formulation of the negative binomial model as previously stated, with the notable change that we model the probability of zeroes for sample *j* and exon *i*, 𝜋, as a function of experimental condition *t*.

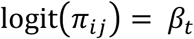

This ZINB model allows for over dispersed counts and additional probability of zeros beyond the expected negative binomial distribution. We use the R package **pscl**’s implementation to estimate parameters^59^. For both the standard and zero-inflated negative binomial approaches, we constructed a corresponding null model by excluding the interaction term 𝛿*_i,t_*, which allows us to compare deviations using a chi-squared test and generate p-values.

3. The third statistical method models the PSI value and total count of reads through a quasibinomial generalized linear model with the experimental condition as a predictor as a more efficient and simplistic model. This allows for using PSI values directly and not having to adjust for exon length in the counts of inclusion or exclusion. Using the quasibinomial formulation, 𝑦*_i_* = 𝑝𝑠𝑖*_i_* ∗ 𝑒*_i_*, where the proportion 𝑝𝑠𝑖*_i_*, the relative use of exon *i*, is equal to the inclusion reads of exon *i* (𝑦*_i_*) divided by the sum of the inclusion and exclusion reads of exon *i (*𝑒*_i_).* Here, we model the reads of exon *I* as:

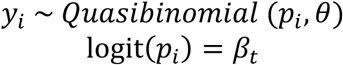

Where 𝛽*_t_* accounts for the impact of experimental condition, 𝜃 is the dispersion parameter, and 𝑝_!_ is the probability of reads covering exon *i*. This method circumvents the exon-length issue and serves as a computationally efficient approach for large-scale evaluations. The generalized linear model was implemented using glm in R with family of quasibinomial. Along with this, as with the negative binomial model we use the standard workflow for dispersion parameter estimation through the function. Here, we extract a p-value using an F-test to compare deviance and test for significance based on impact of experimental condition in nested models. Outlier values are identified using Cooks distance and removed based on a user-specified threshold, with presets thresholds to 4/n (where n is the number of observations/samples) and 1. The default for this outlier detection model is to not be used, however with high numbers of replicates it is useful to extract outliers in the context of the quasibinomial or negative binomial models.

4. Finally, the fourth option is a simple Wilcox Rank-Sum test using PSI values to identify significant shifts across experimental condition. To this effect, the test was performed across experimental conditions *t_1_ and t_2_*. The null hypothesis tested is that the PSI distributions under the two conditions are identical and the alternative hypothesis is that the distributions are not identical:

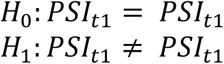

This non-parametric approach provides a robust fourth option for detecting significant relative changes in usage of alternative RNA processing events. We default to applying the Wilcoxon rank-sum test in our processing of differential HIT Indices and changing usage of exon-types across phenotypes. P-values calculated through each method are adjusted for multiple hypotheses to generate a false discovery rate, with a default FDR of 0.05 being considered significant. Alternative RNA processing events are mapped to annotated exons using event-specific Jaccard Indices:

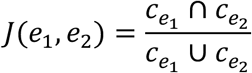

where we are calculating the Jaccard Index between exon 1 (𝑒_1_) and exon 2 (𝑒_2_) and extract the union and intersection between the coordinates of the two exons (*𝑐_e_*_1_, *𝑐_e_*_2_). In terms of exon match selection, SpliceImpactR prioritizes protein-coding, longer, and higher-quality transcripts. We also prioritize similar length transcripts for matching to paired exons to minimize changes outside of the alternative event. We only attempt to match events that occur in terminal exons to exons annotated as terminal. Alternative RNA processing events sourced from the HIT Index require simple matching to individual exons, however, alternative splicing events, such as MXE, require unique search filters such as no exons overlapping a set of coordinates from one transcript that match well in another. This matching process was customized for each alternative event type. Proteins and transcripts associated with each matched annotated exon were retrieved for subsequent analysis. For data sourced from the HIT Index, alternative events—and their associated transcripts and proteins—from the same gene with opposing inclusion preferences were paired for downstream analysis. In cases where more than two events from a gene showed differential inclusion, pairings were made in a pairwise manner between all combinations of upregulated and downregulated events. The data from rMATS is already paired, so no pairing operations are performed on it and the alternative splicing forms are extracted from the coordinates supplied by each event ID.

### Primary sequence local analysis

We assess paired alternative RNA processing events’ transcripts for protein-coding status and classify as such if one or both transcripts are non-protein coding. The amino acid sequences of proteins associated with pairs of protein-coding transcripts containing paired alternative RNA processing events were analyzed for sequence alignment using a BLOSUM62 scoring matrix and an alignment score is extracted to quantify sequence similarity. The alignment score is produced by taking the number of aligned amino acids and dividing by total alignment length. We use the R package msa for this sequence alignment analysis^60^.

Pairs that are both protein coding transcripts are further classified as having a frame shift or not having a frame shift. Frame shifts contain a reading frame shift between the overlapping coding regions in the exons involved in the alternative processing event, leading to the production of different amino acids from the same nucleotides. Frame shifts are identified by comparing reading frames relative to respective transcription start sites (TSS) at overlapping coding regions of each transcript. These overlapping coding regions are restricted to the exons containing the specific alternative splicing event being investigated, however AFE/HFE and ALE/HLE use the closest exons with overlapping coding regions. We identify downstream “rescues” by identifying where the reading frames may realign between the paired transcripts.

The positional distribution of preferentially included exons (proximal vs. distal) is also analyzed within AFE/ALE/HFE/HLE to detect potential global shifts in positional exon inclusion. The pipeline also performs an analysis of protein length, where it compares pairwise protein length. We use a Wilcoxon signed-rank test to identify whether there is a statistically significant shift in lengths between experimental conditions (p-value < 0.05 is considered significant). We also map the distribution of protein length changes in this output.

### Global proteomic analysis

Protein domain annotation was conducted using InterPro to assess domain changes resulting from alternative RNA processing events^37^. We extract InterPro domains by identifying PFAM identifiers through biomaRt^39,40^, then converting to InterPro through the **PFAM.db** R package^61^. InterPro domains removed or added by alternative splicing events were identified (identified by overlap with coding regions of exons), and global domain enrichment analysis was performed to evaluate upregulation or downregulation of specific domains associated with these events. These domains were extracted through identifying when alternative processing impacted sections of the transcript code for sequences of the protein that contain a InterPro domain. If the difference in alternative splicing removes or adds a given domain, it is considered a domain change due to the alternative processing event. A hypergeometric enrichment test was used to identify the globally enriched domains, building background set of changing domains from transcripts matched to the total set of exons identified by the HIT Index. The background set for the enrichment test is produced from the domains identified in the proteins matched to all the exons in the HIT Index’s output (including first, internal, and last exon matches). The p-values from the hypergeometric enrichment test are adjusted for multiple hypothesis testing to calculate a false discovery value, with FDR < 0.05 being considered significant. The tool allows for the removal of repeating domains which change multiple times between one pair of proteins to not overrepresent them in the enrichment plot. We also generate plot output of the distribution of domains changing across each protein pair and the count of proteins pairs with changed domains for each experimental condition comparison.

### Protein-protein interaction changes

Using the domains identified in each protein, a network was built from 3did domain-domain interactions (DDI) to assess when alternative RNA processing events led to changes in the protein-protein interaction networks. Edges were made between isoforms dependent on domain content and DDIs from 3did. We extend a network out one step from each seed isoform node and identify differences in the interacting proteins. These changes were used to identify unique interactions resulting from RNA processing. Extracting the genes containing the novel interactors, SpliceImpactR performs gene set enrichment analysis to understand the functional implications of these novel interactions. We use the R package **hypeR** to generate the geneset enrichment analysis, across Gene Ontology (Biological Process, Molecular Function, Cellular Component), KEGG, and Hallmark genesets^62^. This enrichment analysis defaults to an FDR of 0.05 to identify significance. We also generate output of each network along with noting the novel interactors and interactions.

### Holistic Analysis

To integrate information across multiple alternative RNA processing events, we incorporated a holistic analysis comparing the use of each event. We calculate relative usage of each event by dividing the number of significantly changing events by the total number of events identified in all the samples. We calculate relative domain and PPI changes through the number of domains changed and PPI networks rewired per significant event change. We generate an output table of summary stats for each event covering these statistics, as well as previously generated counts of alignment type (frame shift, rescue, sequence addition, one protein-coding transcript), relative rescue. We also compare alignment scores across event type – identifying which alternative event leads to the largest changes in primary sequence. Along with a summary table, we make various output plots for visualization of these metrics.

### Analyses across human isoforms annotated in the genome of reference

To globally identify transcript swaps due to alternative RNA processing events in the annotated genome, we identified every alternative RNA processing event from the previous 9 categories mentioned, along with alternative transcription start and end sites (aTSS, aTES). aTSS and aTES were defined as overlapping terminal exons with different terminal ends – differentiated from ALE and AFE (and HFE/HLE) which were defined to necessarily require nonoverlap between terminal exons. Using ensemble annotation from **biomaRt**, transcripts were kept if they were part of Ensembl’s transcript support level 1, Ensembl canonical, or Gencode basic transcript sets. Single exon transcripts were filtered out from analysis. Transcripts were also removed if they did not contain complete 3’ and 5’ ends. No isoforms were designated as primary; instead, transcripts were matched pairwise within each gene. Alternative RNA processing events were explicitly identified, with unique requirements for each event type. All exons containing alternative splicing were required to only overlap one exon, except for IR. SE and MXE were not limited to examples with just one exon involved in the exclusive inclusion. Transcript pairs that were both non-protein-coding were filtered out from further analysis. The sets of paired transcripts were used as input for adjusted functionality from SpliceImpactR.

We used the procedure from SpliceImpactR to identify frame shifts and rescues due to annotated RNA processing events. To identify when alternative RNA processing events were co-occurring, we extracted the set of transcript pairs for each event type and used a Dice-Sorensen coefficient to compute similarity metrics in a pairwise manner between the sets:

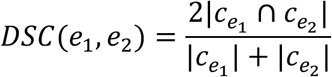

Where we take two times the length of the intersect between the two sets divided by the sum of the lengths of each of the sets. This similarity metric was used because it accounts for large differences in sizes, such as the difference between the number of MXE and the number of aTSS. We also extracted the counts of events that were co-occurring. Looking at how frame shifts and rescues pair alternative RNA processing events, we looked at where the recues occurred and identified what event type the respective exons belong to. We then standardized this by frame shift event count. We used procedures from SpliceImpactR to identify domains and enriched domains. The background set was adjusted to be composed from domains changing across any pair of transcripts from the annotated genome. The enrichment analysis here required the domains to be present in at least 5 genes to ensure one gene with a high number of transcripts wouldn’t be responsible for the domain’s enrichment, given the pairwise approach. We used **dcGOR** to identify enriched gene ontology biological processes associated with the enriched changing domains^63^ (FDR < .05). To identify if similar domains were changing across event type, we used the Dice-Sorensen index to generate a similarity metric between the domains changed across event type in a pairwise manner.

### Analyses across human samples and tissues from Genotype Tissue Expression (GTEx)

Aligned GTEx version 8 RNA-seq data from 17,350 sample across 54 human tissues were downloaded from the AnVIL repository using the gen3-client and checked to be intact with samtools quick-check^64^. The data were then processed with the HIT index pipeline using parameters readnum 10, overlap 15, and bootstrap 5. For added confidence, all hybrid exon calls were required to be assigned a hybrid probability of at least 0.8 by the pipeline’s generative model (see Fiszbein et al., 2022), and AFE and ALE PSI calculations were restricted to exons that were annotated as terminal exons. The aligned data were also processed with rMATS in statoff mode. Using these samples, we used SpliceImpactR on a pairwise basis across each tissue type. We set a threshold of 0.15 for delta psi and 0.05 for FDR. This resulted in 1431 sets of results containing analysis in all 9 alternative RNA processing event types.

We used the **pheatmap** package^65^ in R for each hierarchical clustering with method set as UPGMA. To compare the global changes between tissues, we calculated the average number of paired significantly differentially included alternative RNA processing events across all event types. To identify how the alternative RNA processing events differed in terms of clustering samples, we used the cophenetic correlation between the hierarchical clustering of each of the event types and clustered the resulting correlation. To compare the alternative RNA processing events between blood, brain, and heart, we used the overlap in genes with changing events between tissue type, calculated using a jaccard index, and used UPGMA for hierarchical clustering in pheatmaps’s function. To identify protein domains changing between one tissue and all other tissues, we extracted the list of domains upregulated for the given tissue and assigned all downregulated domains to the other tissue. Filtering out domains which were not commonly changing (with a count below the mean count of each domain), we assigned a value of 1 or 0 for inclusion or exclusion in each tissue type. Using the same UPGWA approach, we performed hierarchical clustering. To calculate relative change in inclusion we took the difference between the mean number of times a domain was present in each tissue and all other tissues.

To identify the relative use of each event type, we first extract the total number of events identified after the previously established filtering for quality of events. We then use the number of significantly changing paired events to generate a relative use metric for each event type. For all geneset enrichment analyses, we used the R package **hypeR** to search for enrichment in various gene sets (Gene ontology, KEGG, Hallmark) using hypergeometric method^62^.

To identify how NLS inclusion is changing due to alternative RNA processing, we used 17 NLS consensus sequences were extracted from Juane et al. (2021).^66^ We then used MEME-suite’s FIMO tool^67^ to scan the amino acid sequences from two sets of proteins for these motifs. The first of these sets is sourced from annotation analysis, all the proteins containing an AFE between them. The second set contains the proteins associated with the AFE that are significantly changing between cerebellum and left-ventricle tissues from GTEx. If within a given pair, only one of the proteins returned a significant match for a NLS motif (q-value < 0.05), this was marked as a change in inclusion of a NLS.

## SUPPLEMENTAL FIGURES

**Figure S1.**
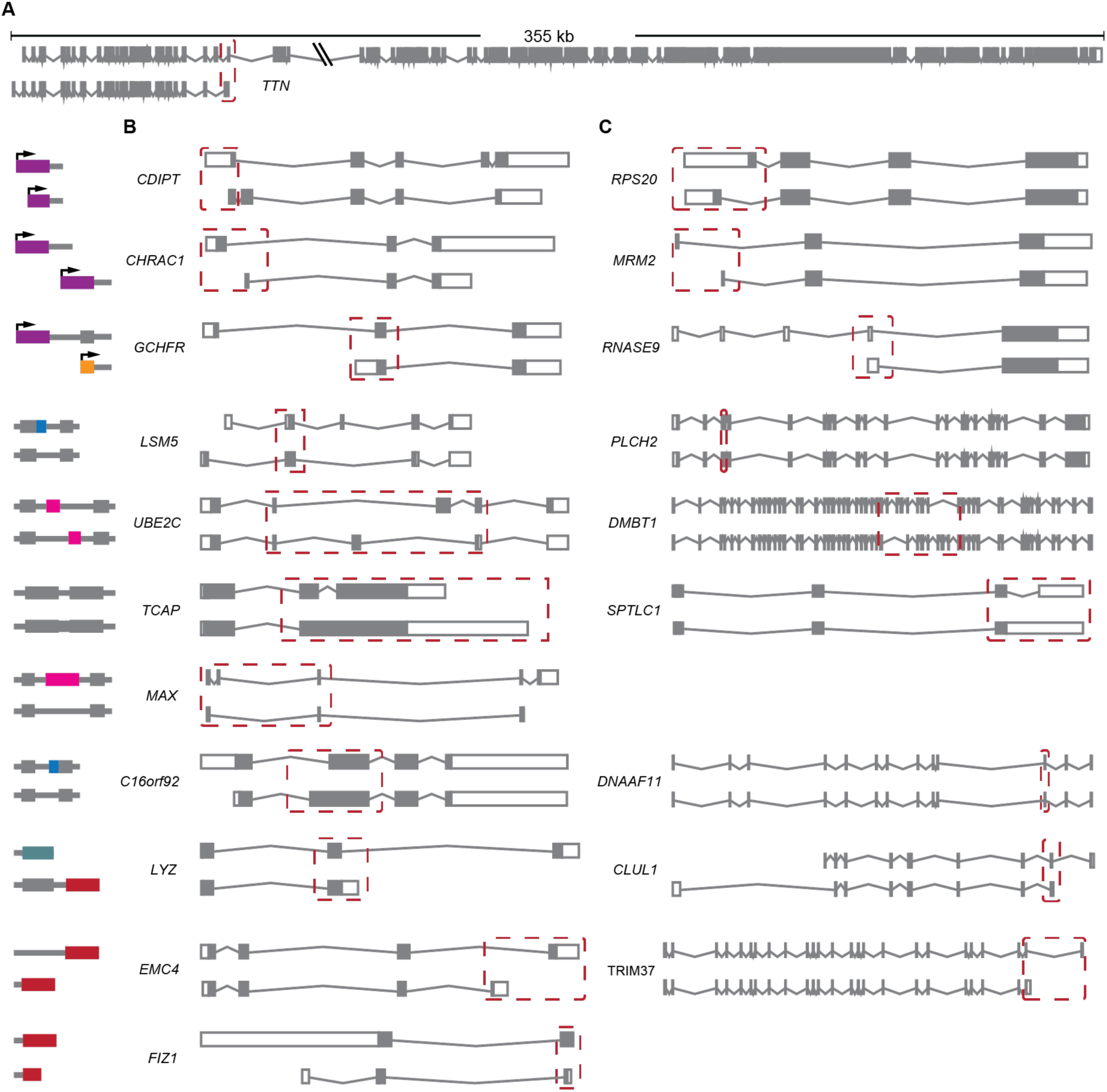
Alternative RNA processing alters transcripts in both expected and unexpected ways, related to Figure 1. (A) Example of an Alternative Last Exon (ALE) event in the TTN gene that alters transcript and protein length by more than 98 kb. (B) Representative transcript isoform pairs for each major type of alternative RNA processing event, where the event occurs within protein-coding regions and results in changes to the final protein product. (C) Representative transcript isoform pairs in which alternative RNA processing events occur within protein-coding regions but do not alter the final protein sequence. No protein-neutral examples were found for aTES. No examples were found that fit this classification for SE or aTES.

**Figure S2.**
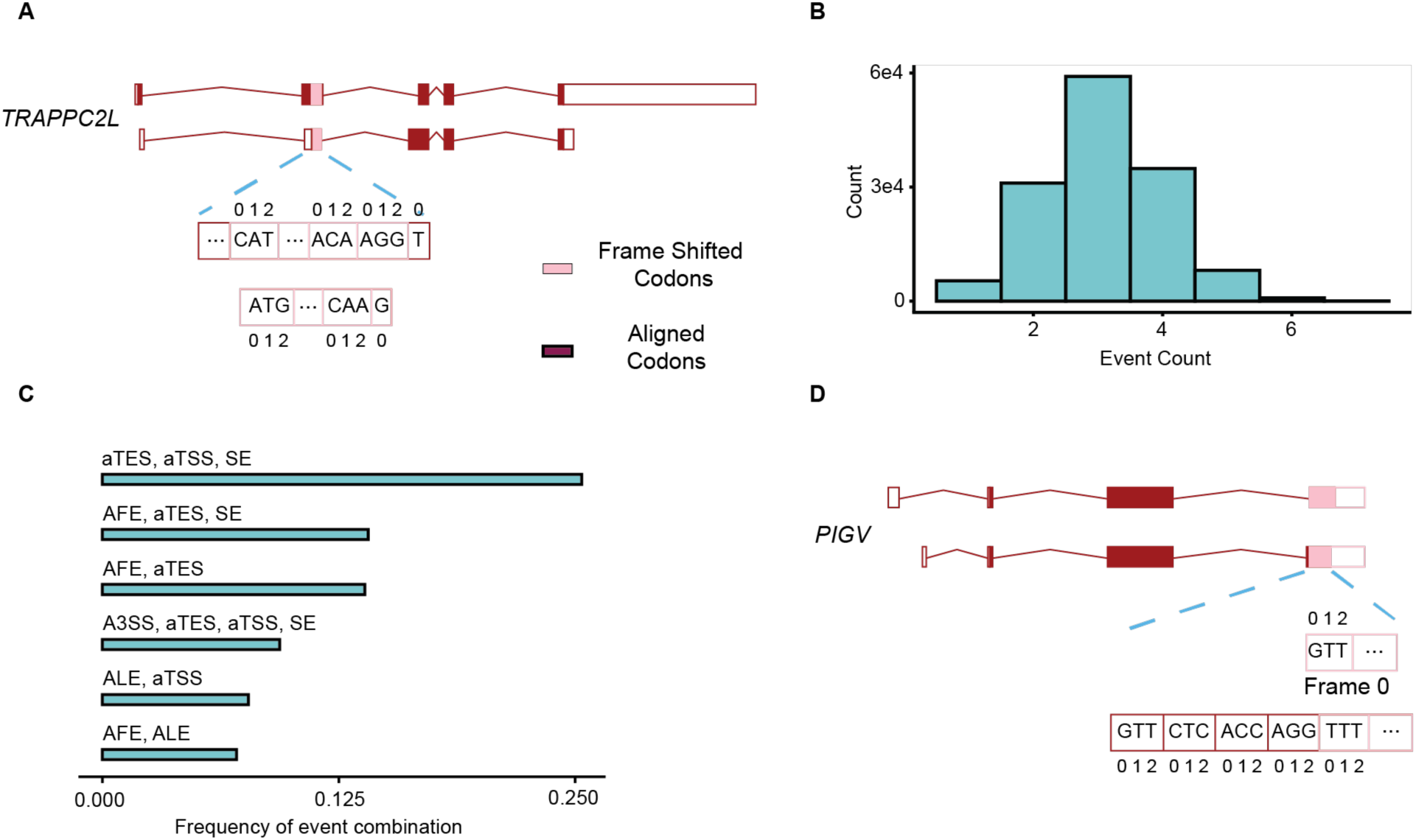
Alternative RNA processing events are co-regulated, related to Figure 2. (A) Example of an in-exon rescue in TRAPPC2L, an A3SS initiates a frame shift and an A5SS rescues the reading frames within the same exon. (B) Count of number of co-occurring alternative RNA processing events per transcript pair when at least one event changes. (C) Frequency of the top co-occurring event combinations across transcript pairs. (D) Example of an A3SS causing a frame shift in just a last exon in PIGV.

**Figure S3.**
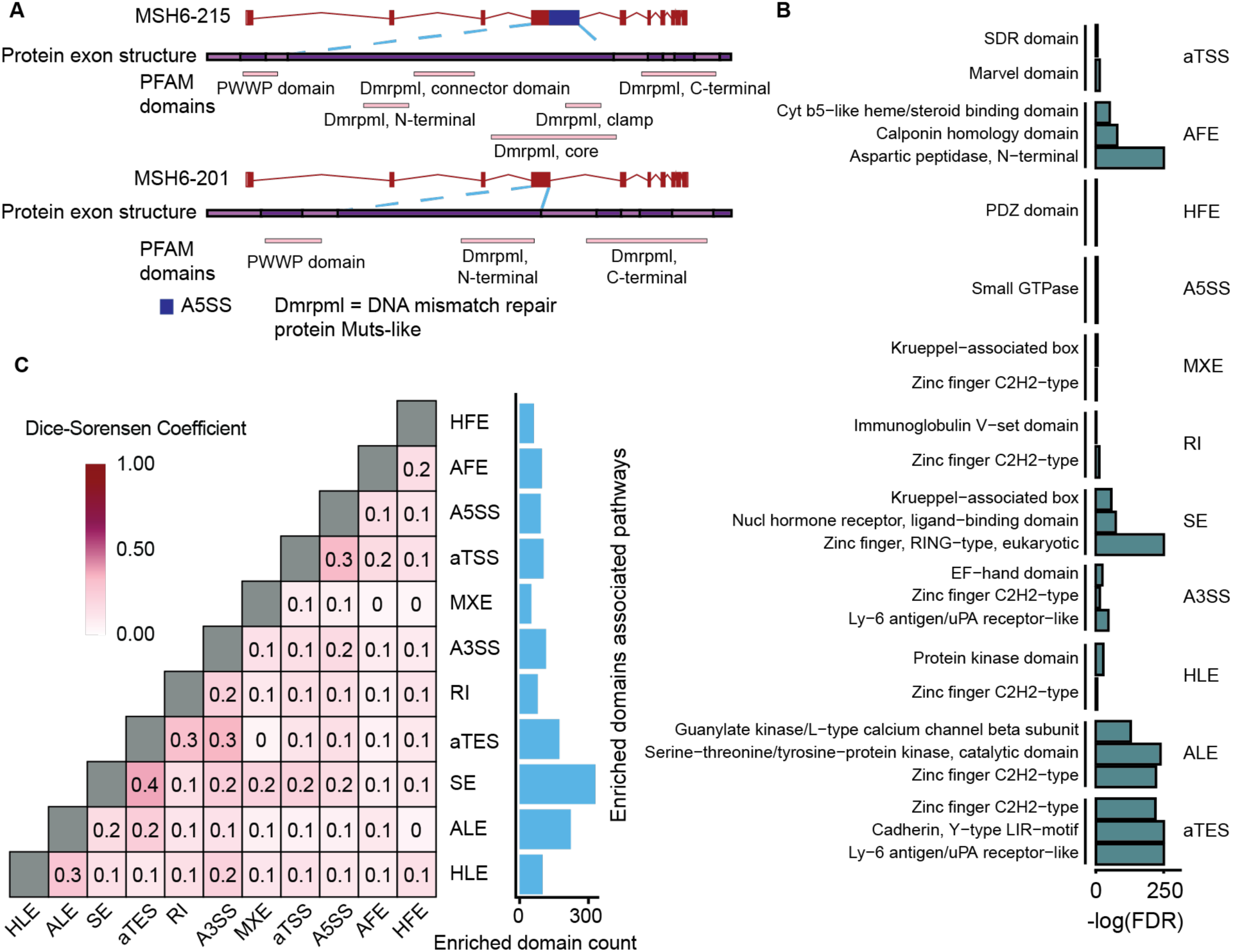
Domain changes due to alternative events drive functional changes, related to Figure 3. (A) Example in MSH6 of multiple DNA mismatch repair protein Muts-like Interpro domains changing due to one A5SS. (B) Enriched domains changing in at least 5 genes due to each alternative RNA processing event. Enrichment calculated through a hypergeometric test relative to a background set of all possible domain changes due to alternative RNA processing. P-values were adjusted to generate false discovery rates (FDR < 0.05 considered significant). (C) (right) Count of enriched domains across each even type and (left) Dice-Sorensen Coefficient overlap between enriched sets of domains from each alternative RNA processing event.

**Figure S4.**
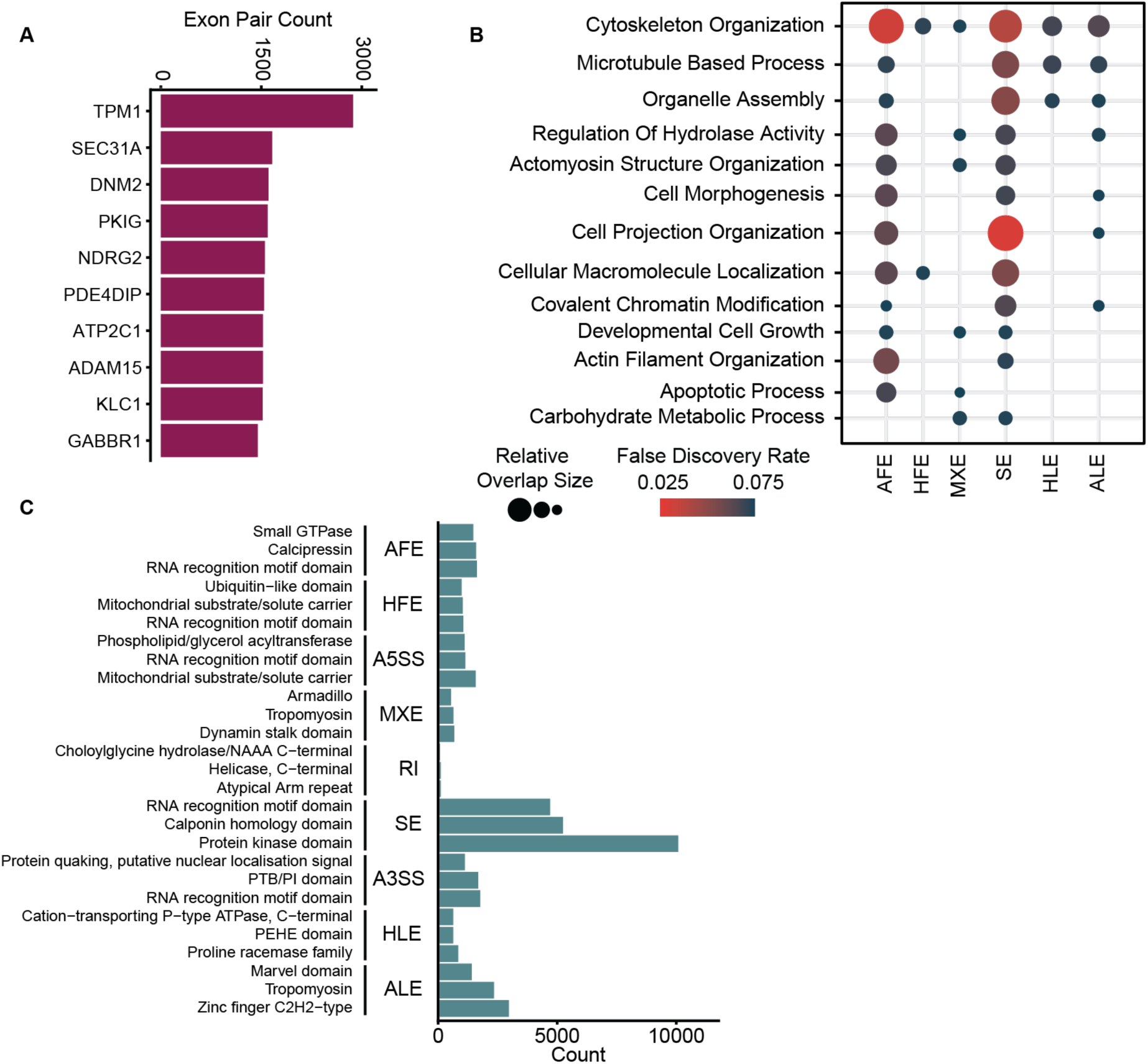
Tissue-specific RNA processing events regulate cell morphogenesis and protein domain inclusion, related to Figure 4. (A) Top 10 genes changing across all tissue and event type comparisons. Exon pair count is calculated through summing the number of tissue comparisons across each alternative RNA processing vent a gene contains significantly changing events. (B) Gene ontology biological process gene set enrichment of genes containing changing alternative RNA processing events across event type. (C) Count of top 3 domains changing due to each alternative RNA processing event.

**Figure S5.**
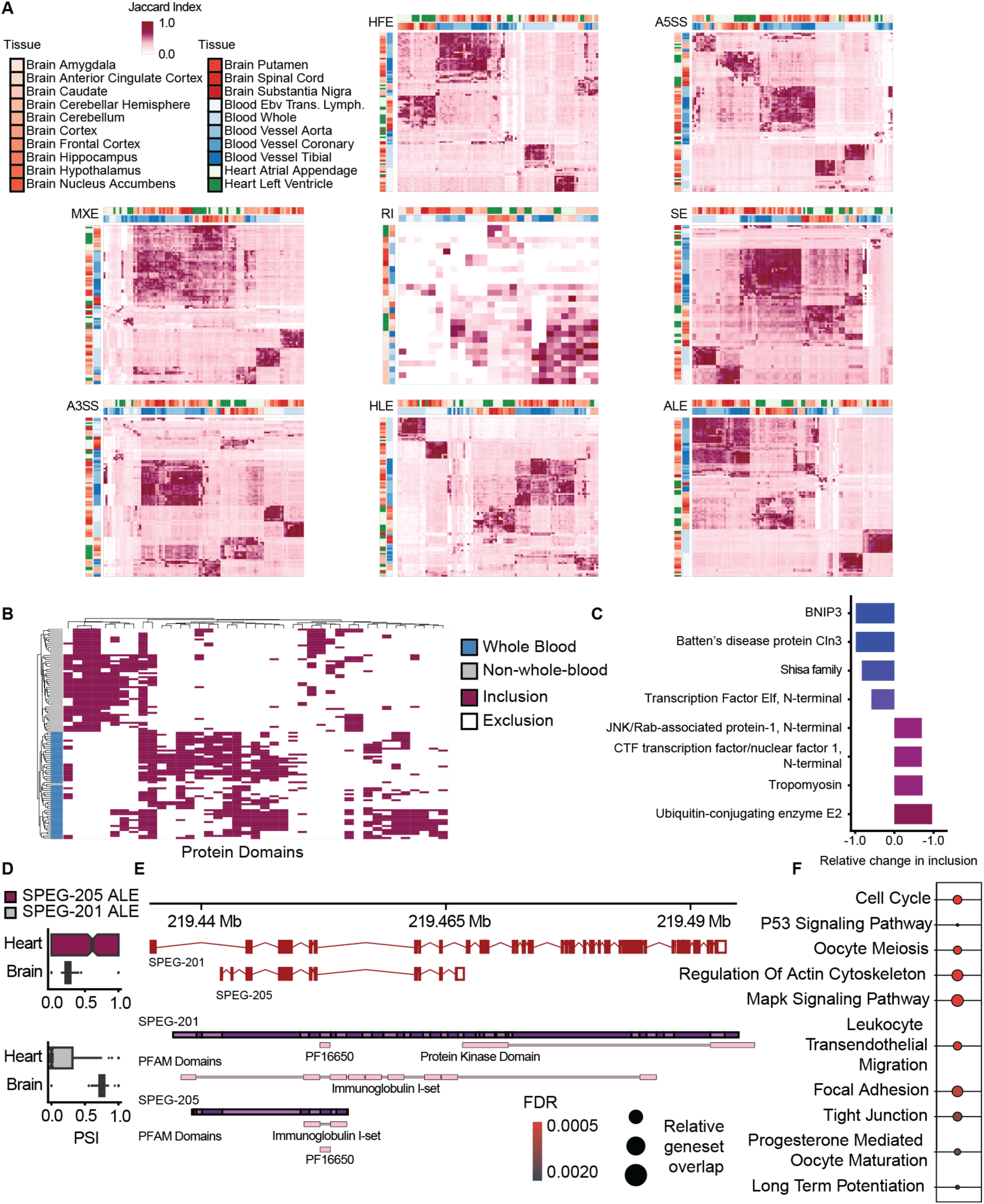
Tissue-specific regulation of alternative events and protein domains, related to Figure 5. (A) Hierarchical clustering using UPGMA on jaccard indices of overlapping genes containing changed alternative events spanning blood, brain, and heart tissues in all alternative RNA processing events other than AFE. (B) Domains upregulated in whole blood were assigned to cerebellum and downregulated to non-whole-blood (presence is inclusion of the domain due to alternative RNA processing). Figure only includes domains present in greater than the median presence number. Hierarchical clustering performed using UPGMA (C) The most regulated protein domains (most included and most excluded) calculated using (count of inclusion / total cerebellum) – (count of exclusion / total non-cerebellum). (D) PSI values of SPEG ALE across cerebellum and left-ventricle. (E) (top) Transcripts matched to ALE identified, (bottom) proteins and domain composition associated with each matched transcript. (F) Gene Ontology Biological Process enrichment of unique PPI due to domain addition in SPEG-201 generated using hypeR.

